# Incipient species or social polymorphism? Diversity in the desert ant *Cataglyphis*

**DOI:** 10.1101/501270

**Authors:** Tali Reiner Brodetzki, Shani Inbar, Pnina Cohen, Serge Aron, Eyal Privman, Abraham Hefetz

## Abstract

Species are the fundamental units upon which evolutionary research is based. In insects, due to the high level of hybridization, the delimitation of such species can be challenging. The genus *Cataglyphis* presents a high level of diversification, making it an excellent model with which to study evolutionary paths. Israel appears to be a “hot spot” for recent speciation in this genus. Although previous studies have described multiple species of *Cataglyphis* in Israel, a recent genetic study has questioned the existence of some of these historically described species. The present study focuses on an apparent species complex that is distinguishable by its mitochondrial DNA (and therefore named mitotypes) but not by its nuclear DNA. Using a multi-method approach (genetics, chemistry and behavior), we show that these mitotypes also differ in their social structures and are readily distinguishable by their cuticular hydrocarbons profiles. While the different mitotypes are in general allopatric, at our study site they all coexist but nonetheless maintain the observed differences between them. This raises many evolutionary questions: Are these incipient species that have diverged with gene flow, or is this a case of social and chemical polymorphism that is maintained within a single species.

## Introduction

Species are the fundamental units upon which evolutionary research is based. Although the biological species concept requires a reproductive barrier between species (Mayr, 2000), many alternative concepts (ecological, evolutionary, cohesion, phylogenetic, and others) that are based on different common characteristics within species, have been proposed (De Queiroz, 2007). Much has also been discussed about whether sympatric speciation can occur and what mechanisms might facilitate it (Bolnick and Fitzpatrick, 2007). Sympatric speciation assumes adaptive divergence of multiple traits in order to generate strong ecologically-based reproductive isolation. However, the possibility of sympatric speciation without some gene flow does not seem probable and, indeed, recent studies have reported that speciation with gene flow is more common than previously thought (Nosil, 2008; Feder, Egan and Nosil, 2012). In insects, incomplete divergence in mating biology can lead to relatively high levels of hybridization between species, challenging the biological species concept and facilitating the possibility of cryptic speciation and speciation with gene flow (Cognate, Seybold and Sperling, 1999; Nosil, Harmon and Seehausen, 2009; Matsubayashi, Ohshima and Nosil, 2010). Generally, speciation can be considered as a continuum on which we can detect various transitions or diversifications such as morphological, genetic, and behavioral, which at different time points may have different levels of differentiation that might not yet show full evolutionary independence, thus named incomplete lineage sorting (Hey and Pinho, 2012; Fišer, Robinson and Malard, 2018). In order to accurately describe diversifications between closely-related species that might be on the species continuum, classical morphology-based taxonomy is not always sufficient, but requires additional methods of identification such as chemical, behavioral, ecological, and molecular (Seifert, 2009; Ross *et al*., 2010; Kather and Martin, 2012; Ronque *et al*., 2016; Fišer, Robinson and Malard, 2018). Differences between species, unlike between populations within a species, should embrace vast changes that cannot be explained as simple polymorphism. For example, cuticular hydrocarbon profiles, the presumed nestmate recognition cues in social insects, slightly differ between nests (exemplifying normal polymorphism) but greatly differ between species (exemplifying major changes). Likewise, behavioral changes that affect either mating preference or mating systems, or foraging, or habitat selection, can all lead to niche displacement and a drive for sympatric speciation (Arnqvist *et al*., 2000). In ants, cases of intranidal mating, gyne (unmated ant queens) morphological polymorphism, a shift from monogyne (single queen) to polygyne (multiple queen) colonies, or host-parasite interactions were all suggested to result in sympatric speciation (Seifert, 2009; Torres *et al*., 2018). This scenario leads to mating place separation and facilitates further genotype divergence, eventually leading to novel genotypes and new species (Seifert, 2010). Social polymorphism (both monogyne and polygyne colonies in the same species and the same population) has been described in several ant species, including *Pheidole, Formica, Leptothorax*, etc. (Bourke and Franks, 1995). Finally, recent studies have shown that there are few cases in which social polymorphism and other traits do not lead to speciation but are acquired by means of a ‘super-gene’, also named ‘social chromosome’ distributed in the population (Wang *et al*., 2013; Purcell *et al*., 2014; Taylor and Campagna, 2016). However, this ‘social chromosome’ mechanism is rather rare and has been described only twice, while most incidents of social polymorphism are mainly believed to depend on ecological constraints.

The ant genus *Cataglyphis* comprises over a hundred species (Agosti, 1990) displaying various social and population structures, ranging from the basic monogynous multicolonial population to polygynous supercolonies, making it an excellent model to study evolutionary paths (Lenoir *et al*., 2009; Boulay *et al*., 2017; Timmermans *et al*., 2010; Leniaud *et al*., 2011; Jowers *et al*., 2013). The genus also expresses various reproductive strategies, from sexual to the asexual (parthenogenetic) reproduction of new reproductive queens (Leniaud *et al*., 2012; Eyer *et al*., 2013; Darras, Kuhn and Aron, 2014; Aron, Mardulyn and Leniaud, 2016). Israel appears to be a “hot spot” for recent speciation in this genus: in the *bicolor* group, we identified eight mitochondrial types (Eyer *et al*., 2017), in the *albicans* group there are nine identified mitochondrial types (Eyer and Hefetz, 2018). These mitochondrial types do not always concur with nuclear delimitation methods, and it is not clear whether mitochondrial types that have been previously used to delimit species can also serve as identifiers in this case of the *bicolor* group.

Here, we examined an apparent species complex of the previously described putative species *C. niger* and *C. savignyi* (Agosti, 1990; Radchenko, 2001), and a population tentatively identified as *C. drusus* (henceforth termed the “niger complex”). In a previous study, these putative species were found to exhibit different mitochondrial haplotypes (hereafter called mitotypes), but could not be differentiated on the basis of the nuclear genes used (Eyer *et al*., 2017). To disentangle this species complex, we used further genetic tools (ddRAD sequencing and microsatellite markers), together with the chemistry of cuticular hydrocarbons, and behavioral assays designed to examine colony insularity.

## Methods

### Sample collection

Sample collection was divided according to the different analyses methods: 1. Samples collected from different geographic areas across Israel for the RAD sequencing analysis 2. samples collected from the Betzet and the Tel-Baruch populations for the microsatellite and the CHC analyses; 3. samples collected from the Betzet and the Tel-Baruch populations for behavioral experiments; and 4. samples of queens, newly mated gynes, and their spermatheca contents for mitotype analysis from the Tel-Baruch population.

1. 118 individual were collected from 55 locations across Israel, and from Turkey (kindly collected by Dr. Kadri Kiran, Trakya University, Turkey), 76 were originally collected along with samples for Eyer et al. 2017 and the rest 42 samples were newly collected for this project (sample collections are detailed in S1). All samples for genetic analysis were kept in absolute ethanol until extraction.
2. Nests were collected from two coastlines locations: Betzet (33° 4’36.09”N, 35° 6’32.05”E) and Tel Baruch (32° 7’33.76”N, 34°47’9.04”E). These locations present similar ecological niches that are characterized by semi-stabilized sand dunes. We sampled all visible nests in a plot of 200*200 m, as well as a transect of 1 km (at 100 m intervals). From the Betzet and Tel Baruch sites respectively, we collected partial nests-queenless nests (QL; n= 57/39), and whole nests/queenright nests (QR; n= 2/11; at). All the nests in the experimental plot were excavated over four consecutive days to avoid pseudo-replication due to nest migration. Ants were dissected as follows: the head, for CHC content, was kept in hexane; the thorax and legs, for DNA extraction, were kept in 100% ethanol; and the abdomen was frozen at -20°c and kept for future analysis. Table S2 details the number of nests used in each analysis and collection site.
3. Nests at the above-mentioned locations were thoroughly excavated to include the queen, all workers, and brood. To ensure that the entire nest was collected, excavation was further extended by an extra 0.5–1 m in all directions after having observed the last gallery. In the Betzet location, despite our thorough excavations the queens were difficult to locate, so we used both QR (n=3) and QL nests (n=15) for the experiments. At Tel Baruch we successfully excavated and used 10 QR nests.
4. Established queens were collected from the above excavated nests. Newly mated gynes were collected from the Tel Baruch population during the 21 consecutive days of nuptial flight. Some of the gynes had shed their wings and some had also begun to dig a new nest. All were dissected and their spermatheca content was extracted. Sperm was kept in Ringer’s buffer and frozen at -80°c until DNA extraction.

### Genetic analysis

#### Mitochondrial haplotypes

were determined by genotyping Cytochrom B using one worker per nest (Eyer et al., 2017). CytB primers used: CB1 (Forward) TATGTACTACCATGAGGACAAATATC and CB2 (Reverse) ATTACACCTCCTAATTTATTAGGAAT, annealing temperature was 48°C (Simon *et al*., 1994). PCR products were purified with ExoSAP-IT™ PCR purification kit (Applied Biosystems). Sequencing was performed on an ABI 3730 Genetic Analyzer (Applied Biosystems). Base calling and sequence reconciliation were inspected by eye and performed using Unipro UGENE v 1.3. Sequences were aligned using MUSCLE algorithm (Edgar, 2004) implemented in UGENE Aligner, together with sequences previously used in Eyer 2017.

#### RAD sequencing

We held suspicions that the genetic markers employed in Eyer *et al*. 2017 were possibly not sensitive enough to detect differences between the mitotypes so we decided to use Double Digest Restriction Associated DNA (**ddRAD**) Sequencing that provides many more genomic markers enabling better accuracy for delimiting species. All the DNA for this analysis was extracted using Qiagen blood and tissue DNA extraction kit following the standard protocol. We used a modified protocol of (Parchman *et al*., 2012; Peterson *et al*., 2012), to construct double-digest RAD libraries, modifications added by A. Brelsford, A. Mastretta-Yanes, J. Leuenberger, R. Sermier. Changes were as follows: Added dual-index barcoding to allow multiplexing >96 samples per library. The Y-adapter for MseI from Peterson et al (ddRAD) was used to prevent amplification of Msel-Msel fragments. Simplified restriction and ligation mixes using CutSmart buffer. Purification post-ligation was done using Agencourt AMPure beads. PCR using Q5 Hot Start Polymerase (New England BioLabs).PCR modified to decrease number of cycles by increasing number of replicates and starting DNA volume. Addition of primers and dNTPs for a final thermal cycle, in order to reduce production of single-stranded or heteroduplex PCR products. Final cleanup and size selection was carried out with AMPure beads. Three samples were omitted from the final results due to poor sequencing yield. All samples were multiplexed and sequenced by single-end 100bp reads on 2 lanes of an Illumina HiSeq4000 sequencer.

Processing of RAD sequence data- A total of 648,303,541 reads were sequenced, with an average of 5,026,520 reads per sample (275,586 reads for the lowest coverage sample, pol633_u). Dividing the number of reads by the 67,687 EcoRI restriction sites in the reference genome, it is an average coverage of 37X. The raw reads were initially processed using the *Stacks* pipeline (Catchen *et al*., 2011; Catchen, 2013) and low quality reads were discarded. These were defined as reads in which the score drops below an averaged phred score of 10 in a sliding window of 15% of the reads’ length, that is 14 bases for a read length of 93 bases. After this initial processing, 583,275,943 reads that were mapped to a sample remain (92%). A total of 459,081,529 reads were mapped to the reference genome of *C. niger* (version Cnig_gn1) using *Bowtie2* (Alexander, Novembre and Lange, 2009). We filtered the alignments allowing for a maximum of four mismatches. To filter non-unique mappings, we removed alignments where the second best hit had less than twice the number of mismatches found in the best hit. After this filtering, 369,719,665 alignments remained. At this point, the actual coverage of mapped loci was calculated, with an average of 22X. We continued to analyze the mapped sequences to identify single nucleotide polymorphic sites (SNPs) using the *Stacks* pipeline and created a catalog containing 300,509 SNPs for the 126 samples.

*STRUCTURE* analysis for RAD samples-Individual samples were divided to seven populations according to their mitotype: *israelensis, drusus, niger, savignyi, holgerseni, isis* from Israel, and from Turkey-*noda*. The data in the catalogue was filtered as follows: 1) loci were removed if they had more than 7% missing data (samples where a genotype could not be called at that locus). 2) Minimal minor allele frequency (MAF) of 1% (a minimum of 3 allele copies in the 230 chromosomes of 115 diploid individuals). 3) A minimum of five reads per individual for calling a genotype. Each sample was required to have genotype calls in at least 80% of the loci. This resulted in a catalogue of 19,995 high confidence SNPs that were used in the population structure analysis (supplementary Fig. S1).

*STRUCTURE* (Pritchard, Stephens and Donnelly, 2000; Falush, Stephens and Pritchard, 2007) was run over *K* (number of expected clusters) values of 3 – 10; for each *K* value, 100 iterations of *STRUCTURE* were run, each over a different set of 1,000 SNPs randomly chosen out of the high confidence SNPs. In each set, a minimal genomic distance of 2,000 nucleotides between sites was maintained to comply with STRUCTURE’s assumption of no linkage disequilibrium. The program was run with 1,100,000 MCMC iterations, and the first 100,000 were discarded (burn-in period). All the other parameters were kept at default. The results were analyzed using *CLUMPAK* (Kopelman *et al*., 2015) and combined for each K value.

To test whether the three species in the complex are genetically separated or not we used full maximum likelihood calculations to compare between two multi-species coalescent models as implemented by 3s program (Yang, 2002; Zhu and Yang, 2012; Dalquen, Zhu and Yang, 2017). The null model which does not allow gene flow between two closely related species and an alternative model that does. Both models include a third species that is used as an out-group. This species is assumed no to have genetic flow with the species tested. We preformed multiple such likelihood ratio tests (LRT) between different combinations of individuals of the populations of *C. savignyi, C. drusus, C. niger, C. israelensi* and *C. isis*, first to check for gene flow between *C. israelensis* and any one of the species in the complex with the out-group of *C.isis* and then between the species of the complex with the out-group of *C. israelensis*.

93 bases long sequences of these individuals, product of the RAD sequencing, were aligned to a reference genome of *C. drusus* as before, (allowing a maximum of 4 mismatches for the best hit and a minimum of twice as many mismatches as the best hit for the second best hit), only this time no gaps were allowed. Based on the position of the sequences in relation to the reference genome, 3 sequences from 3 different individuals were aligned together in one of the following configuration: 123 – a sequence from individual of species 1, a sequence from an individual of species 2 and a sequence from individual of species 3; 223 – two sequences from two different individuals of species 2 and a sequence from individual of species 3; and similarly 113. This is assuming species 1 and 2 are tested for gene flow while species 3 is used as an out-group. At least 2000 bases gap was maintained between the triplets of aligned sequences (as inferred from the reference genome). For each locus one of the three possible alignment configurations was chosen at random.

For each combination of species, 3s was run three times over the corresponding aligned sequences, using the Gaussian quadrature number of points = 16 and different seed values. Then LRT was performed between the highest scoring null and alternative models. The null model of no genetic flow was rejected if 2DlnL > 5.99 (corresponding to ***χ***^2^ distribution with two degrees of freedom and significance level of 0.05 for type I mistake).

#### Microsatellites

To understand the genetic population structure as well as the social structure we used a microsatellite markers analysis. The analysis was performed on 648 workers of the Betzet population *(C. drusus* only; X+SE=11.42+0.1, N=57 nests). DNA was extracted with 5% CHELEX (BIO-RAD) and then amplified with 11 microsatellite markers (previously designed for: *C. hispanica* - Ch01, Ch08, Ch10, Ch11, Ch12; Darras, Kuhn and Aron, 2014, *C. cursor* - Cc54, Cc65, Cc96 and Cc11; Pearcy *et al*., 2004, and *C. niger* - Cn02, Cn04; Saar *et al*., 2014). Similarly, genetic analysis was performed on 548 workers of the Tel Baruch population (X+SE=21.08+0.45, N=26 nests). DNA was amplified with 7 microsatellite markers (previously designed for *C. hispanica* - Ch23; Darras et al. 2014, *C. cursor* - Cc99, Cc54, Cc51; Pearcy et al. 2004, and *C. niger* - Cn02, Cn04, Cn08; Saar et al. 2014). Amplification was done using Type-it PCR mix (QIAGEN). PCR products were sequenced with ABI3500, genotypes were then analyzed with GeneMarker. Microsatellite data were checked for linkage disequilibrium, fixation index, and G-test using GENEPOP (Raymond and Rousset, 1995), heterozygosity was checked using both GENEPOP and FSTAT (Goudet, 1995), and F_st_ was checked using FSTAT. To assess the number of queens, queen genotypes, and the number of matings we used the program COLONY (Jones and Wang, 2010). The program STRUCTURE (Pritchard, Stephens and Donnelly, 2000) was used to evaluate the number of colonies in the population. The number of K (number of possible colonies, from 1 to n size) was evaluated by the Evanno method using STRUCTURE Harvester (Earl, 2012). PCA of microsatellite data was performed as described by Ryan *et al*., 2017 (using R version 3.3.3; library *adegenet* and *ggplot2* for visualization), and relatedness was estimated using Coancestry (Wang, 2011; Queller GT method).

### Behavioral assays

In order to examine nest insularity and colony composition at the population level we used nest-merging experiments (Reiner Brodetzki & Hefetz 2018). We composed queenright (QR) colonies (n=3 and n=10 for Betzet and Tel Baruch, respectively) comprising approximately 200 workers maintained in nesting boxes prior to the experiment, as well as freshly excavated queenless (QL) nests (n=15). Workers and queens of each colony were marked on the thorax with different colors of nail polish, and then left for a day to get acclimate. At the onset of the merging experiment the two nests were connected to a common foraging arena (size 20×20 cm). Sand and food were placed in the foraging arena and water and cotton balls in the enclosed nests. Scan observations were performed for up to three days (merging was stopped on two occasions in order to save the queens), while recording interactions between non-nestmates, including aggression, trophallaxis, adult carrying, and antennation. For the Betzet population, merging experiments were carried out between 18 nests, three of which were QR and 15 were QL. The merging tests between the QL nests were performed up to a week after excavation, in order to avoid a possible ‘QL nest behavior’, in which nests that are rendered queenless have a tendency to lose their specific nest ‘signature’ over time. All nests were tested against all nests. If nests merged together, they were left as one colony. If we observed over ten aggressive interactions within ten minutes, we stopped the test and separated the two nests. In Tel Baruch merging was done using 10 QR nests of the different mitotypes, as detailed in Table 4. If nests merged, they were left together as one colony with multiple queens.

### Cuticular hydrocarbon analysis

In order to distinguish colony boundaries according to their cuticular hydrocarbon profiles, we immersed individual heads in hexane (5–10 ants from each nest), containing 100 ng/μl of eicosane (C_20_) as internal standard. Initial analysis was conducted by gas chromatography/mass spectrometry (GC/MS), using a VF-5ms capillary column that was temperature-programmed from 60°C to 300°C (with 1 min initial hold) at a rate of 10°C per min, with a final hold of 15 min. Compound identification was achieved according to their fragmentation pattern and respective retention indices compared to authentic compounds. We identified 74 and 60 compounds for the Tel Baruch and Betzet population samples respectively (supplementary Fig. S2 A+B). Quantitative analyses were performed by flame ionization gas chromatography (GC/FID), using the above running conditions. Peak integration was performed using the program Galaxie Varian 1.9. A total of 37/35 hydrocarbons where used in the Tel Baruch and Betzet analysis respectively (highlighted in bold in supplementary Fig. S2 A+B). Hydrocarbon profile specificity was determined using multivariate statistics, i.e., discriminant analysis using the stepwise forward mode. This module of the program works initially as a principle component analysis for reducing the number of variables to fit the number of cases, and then performs the discriminant function. We then used k-means clustering in order to determine the structuring of the population, and assessed the most probable K using kmeans function in *stats*, as well as mclust in *mclust* and nbclust in *NbClust* libraries in R. Plot was done by clusplot function in *cluster* library.

## Results

### RAD sequencing

In order to clarify further the taxonomic status and species delimitation of the *bicolor* group in Israel (see Eyer *et al*., 2017) we performed RAD sequencing on samples from different localities, which were further divided into groups based on their mitotype *C. drusus* (n=9), *C. niger* (n=14), *C. savignyi* (n=47), *C. israelensis* (n=35), *C. isis* (n=4), *C. holgerseni* (n=3), and samples from Turkey (n=6) identified as *C. nodus*.

*STRUCTURE* analysis for the possible K clusters was examined between 3 to 10 numbers of K’s, resulting in the most probable Ks of 5, 6, and 7 (Figure 1). There was a clear separation between *C. isis, C. holgerseni* and *C. israelensis* at the 3 possible Ks, and in k=7 also *C. nodus* was clearly distinct. However, at K=6 or 7) distinction between the *C. holgerseni* and *C. nodus* samples was not clear cut (green or red and green respectively). In the *C. israelensis* population, some individuals showed genetic polymorphism that suggests hybridization between *C. israelensis* and the *C. niger complex*, mainly within the *savignyi* mitotype. *STRUCTURE* analysis for any attempted K (between 3 to 10) could not separate the three mitotypes populations of *C. drusus, C. niger* and *C. savignyi* and they are all clustered together (Fig. 1; colored in orange).

**Figure 1:**
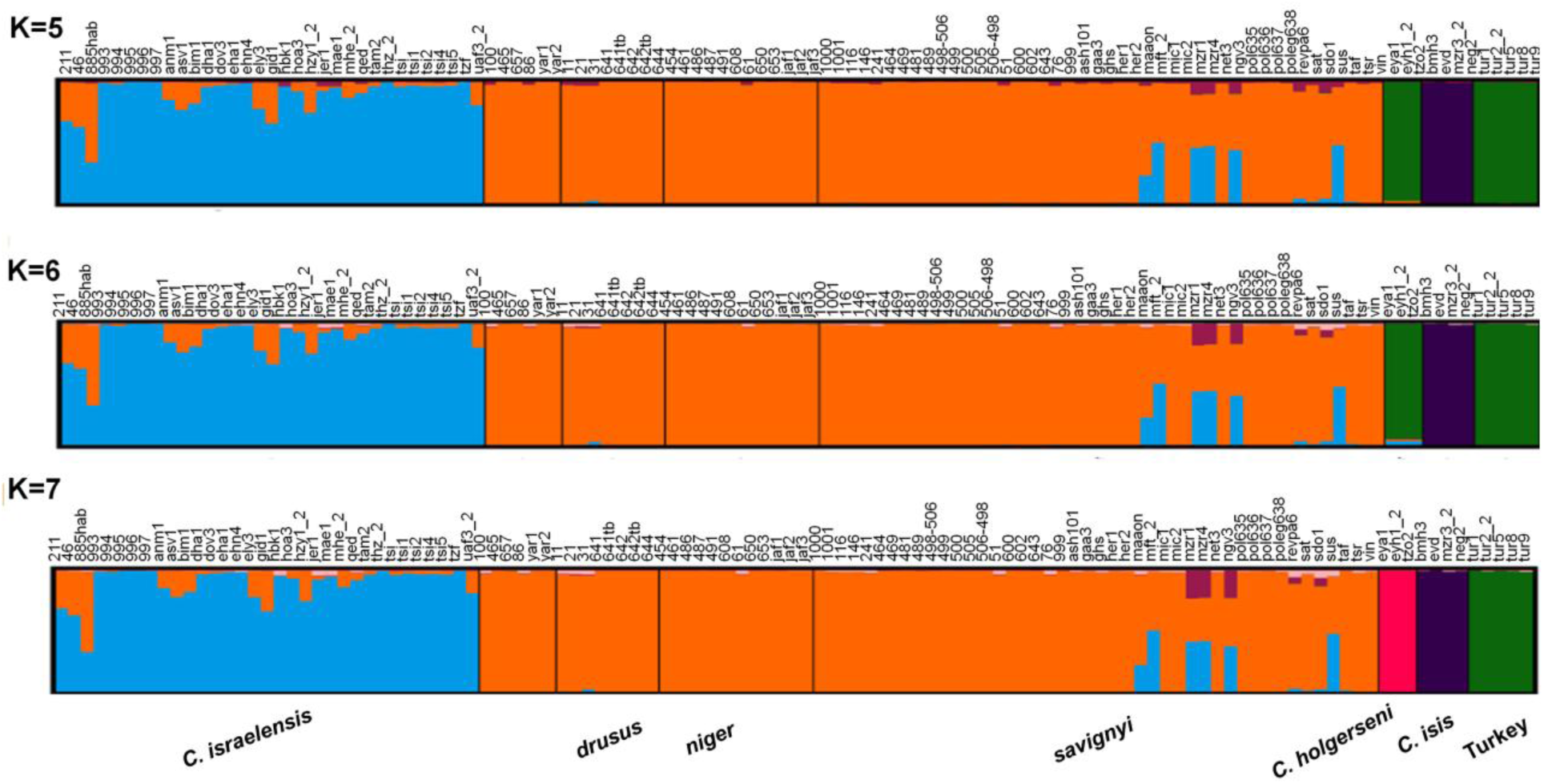
STRUCTURE results of the ddRADseq, possible K outcomes 5-7. *C. israelensis* in light blue, the *C. niger* complex: *C. drusus, C. niger*, and *C. savignyi* in orange, *C. holgerseni* in green or pink, *C. isis* in dark blue, and *C. noda* in green.

Species delimitations within the *niger* complex were put to the test using the maximum likelihood approach as implemented by 3s program, comparing a coalescent null model that does not allow for gene flow between tested species and an alternative model that does. At first *C. israelensis* was tested against each of the species in the complex with the out-group of *C. isis*. As the null model could not be rejected for any of the tests, we used *C. israelensis* as the out-group in the following tests to check for gene flow within the *niger* complex. In all three tests the null model was rejected and the *savignyi, niger* and *drusus* mitotypes were found to have gene flow between them. For the combination of *C. Holgerseni, C. savignyi* and *C. isis, C. Holgerseni* was found to have gene flow with the *savignyi* mitotype (Table 1).

**Table 1:** Gene flow 3S test. 3S run results comparing two coalescent models for different triplets of species. The difference shown is between the highest scoring alternative model and the highest scoring null model between 3 runs for each of the species combination. The bold lines are for species triplets, for which the null model was rejected, suggesting gene flow between the first two species.

| Species combination | loci Nr. | 2DlnL | pvalue |
| --- | --- | --- | --- |
| *C. israelensis* : *C. savignyi*, *C. isis* | 23,031 | 0 | >0.99 |
| *C. israelensis* : *C. niger*, *C. isis* | 22,770 | 4.933 | 0.1 |
| *C. israelensis* : *C. drusus*, *C. isis* | 23,662 | 0.55 | 0.975 |
| *C. savignyi* : *C. niger*, *C. israelensis* | 27,769 | 103.99 | **<0.001** |
| *C. savignyi* : *C. drusus*, *C. israelensis* | 28,830 | 384.18 | **<0.001** |
| *C. niger* : *C. drusus*, *C. israelensis* | 28,626 | 57.399 | **<0.001** |
| *C. Holgerseni* : *C. savignyi*, *C. isis* | 22,771 | 21.17 | **<0.001** |
| *C. savignyi* : *C. isis*, *C. holgerseni* | 23,561 | 0.008 | >0.99 |

### The Betzet population

#### Genetic analysis

We examined a subset of 25 nests from the Betzet population for their mitotypes, and found that all of them presented the same mitotype, similar to the *drusus* group described in Eyer et al. (2017).

Microsatellite DNA analyses revealed that out of the 11 markers amplified and sequenced, three (Cc65, Cc96 and Cc11) were at linkage disequilibrium and therefore omitted from the analysis. The number of alleles in the remaining eight microsatellites loci ranged from 4 to 16. Mean heterozygosity was *Ho=* 0.65 (range 0.31–0.84). The fixation index *Fit* was slightly higher and significantly different from zero (mean ± SE jackknife= 0.09 + 0.06; Z test= 11 p-value< 0.0001), indicating deviation from Hardy-Weinberg equilibrium and suggesting that mating was not completely random in the population studied. *Fst* across the 59 nests sampled at Betzet was 0.19, while within nest relatedness was 0.35 (SE jackknife: 0.02). The G-test showed a result significantly different from zero between some of the nests, indicating that they were genetically differentiated and belonged to distinct colonies (21 nests), while for other nests the G test was not significantly different from zero and they are considered to belong to the same colonies (36 nests divided into 13 colonies). STRUCTURE analysis indicated that the population consists in 2, 3, or 13 genetic colonies (most probable K numbers; Fig. 2 A-B) out of 57 nests analyzed. However, this result of 13 colonies did not seem very probable due to the dispersed location within the plot of the nests assigned to a particular genetic colony, and the fact that mixed genotypes were present within a single nest. K= 2 did not seem very probable either, as it suggests a unicolonial population (we did not see any evidence for such a population structure when observing the population in the field). We therefore conducted a Principle Component Analysis using the genetic colonies assigned by STRUCTURE division of K= 3, which revealed only a very small differentiation in genotypes, which were mainly very similar (PC1= 3.4%, PC2= 3.1%; Fig. 2C).

**Figure 2:**
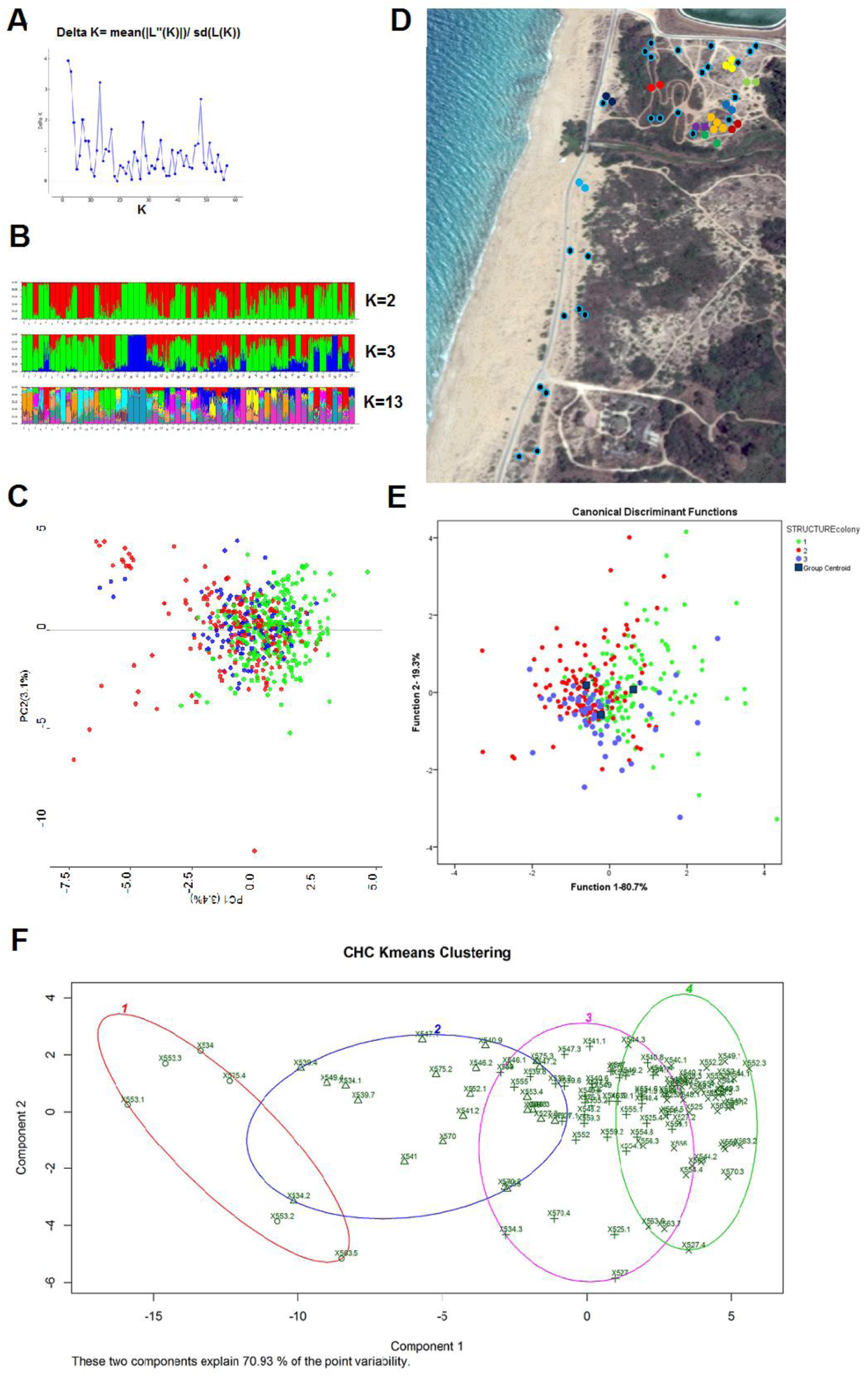
A) STRUCTURE harvester results for the Betzet population indicating the best probable K is K=2, then K= 3 and K=13. B) The population STRUCTURE divisions according to the most probable K possibilities. C) Principle Components Analysis of genotypes colored according to STRUCTURE result k=3 D) Locations of estimated colonies based on similar assumed matrilines according to COLONY; same color indicates the same matriline, light blue circle with black edge indicates a colony with no other similar matriline. E) Discriminant Analysis of CHC groups are set according to k=3. F) K-means of CHC profiles K=4, individuals are marked with nest number (first 3 digits) and individual ID. Circle indicates k-1, triangle k-2, cross k-3, and X k-4. The two components explain 70.93% of the variation.

Genotyping of workers from 42 nests showed that within each nest all of the workers’ genotypes matched to a single queen; this conclusion was also supported by COLONY, unequivocally confirming monogyny. In the remaining 15 nests some of the worker genotypes (average of 1.7 workers per nest; SE= 0.22) did not perfectly match the queen’s assigned genotype, suggesting either multiple queens (which is possible but less likely in light of the above data indicating monogyny), queen turn-over with workers that had remained in the nest from previous residing queens or the presence of drifting workers (workers who mistakenly enter and remain in another colony; a phenomenon documented in other *Cataglyphis* species; M. Knaden, personal communications). Comparing the reconstructed matrilines of different nests based on the COLONY results, we assumed that nests with similar matrilines belong to the same colony, resulting in several polydomous colonies found in the population (Table S3 and in Fig. 2D). Considering monogyny, the COLONY analysis suggested polyandry with an average of 5.3 (SE= 0.22) males contributing to worker production (range: 2-9 males).

#### Cuticular hydrocarbons

Cuticular washes contained 60 identified hydrocarbons, of which 35 were used in the following analyses (supplementary Fig. S2A). CHC profiles were analyzed by calculating the relative amounts of each compound (peaks comprising inseparable compounds were considered as a single compound), for 49 nests (275 individuals). We then used the k-means approach to determine the level of structuring in the samples. The most probable number of K’s was 4 according to all three analyses methods used (k means, mclust, and nbclust), with the two components explaining 70.93% of the variability (Fig. 2F). We further performed a discriminant analysis based on the CHC profiles of the genetic colonies assigned by ‘STRUCTURE’ at K= 3. Colony assignment of three explains a high percentage of the variance (function1-80.7% and function 2–19.3%), although the profiles are very similar and overlap (Fig. 2E).

#### Behavior

A possible resolution of the presumed discrepancy between CHC and genetic data lies in performing behavioral assays, that is, in examining whether two distinct nests will merge to a single colony when given the opportunity. Out of the various 27 trials performed, 14 nest-pairs successfully merged, culminating in seven colonies, each encompassing one to five nests (Table 2 and Fig. 3). During the successful merges there was no aggression at all, either between workers or between workers and queens. The remaining 13 trials resulted in mutual aggression between workers that belonged to different nests, as well as aggression towards the queens (in QR nests). These trials were stopped after ten minutes to avoid further injuries to workers and queens. When merging occurred it was between either a QR and a QL nest or two QL nests, and generally when nests were located in close proximity (except for one case), suggesting that colonies with the *C. drusus* mitotype in the Betzet population are generally polydomous, having one central QR nest and several satellite QL nests.

**Table 2:** Merging assays of the Betzet population

| Nest A | Nest B | Merging | Between-colony trophallaxis | Between-colony aggression | Maximal distance between colonies (meters) |
| --- | --- | --- | --- | --- | --- |
| 1 | 9 | + | + | - | 1.5 |
| 2 | 3 | + | + | - | 2 |
| 4 | 5 | + | + | - | 1.5 |
| 14 | 15 | + | + | + | 2 |
| 10 | 11 | + | + | + | 2.3 |
| 16 | 17 | + | + | + | 10.4 |
| 6 | 4+5 | + | + | + | 4.5 |
| 8 | 13 | - | - | + | 14 |
| 12 | 13 | - | - | + | 20 |
| 8 | 12 | + | + | - | 18 |
| 2+3 | 4+5+6 | - | - | + | 34 |
| 1+9 | 4+5+6 | + | + | - | 8.5 |
| 1+9 | 2+3 | - | - | + | 10 |
| 16+17 | 14+15 | - | - | + | 37 |
| 10+11 | 2+3 | - | - | + | 36 |
| 16+17 | 18 | + | + | - | 9.4 |
| 14+15 | 12+8 | - | - | + | 15-50 |
| 1+9+4+5+6 | 10+11 | - | - | + | 20 |
| 1+9+4+5+6 | 12+8 | - | - | + | 37 |
| 1+9+4+5+6 | 7 | + | + | - | 22 |
| 10+11 | 12+8 | + | + | - | 55 |
| 14+15 | 13 | + | + | - | 42 |
| 14+15 | 10+11 | - | - | + | 75-112 |
| **10+11** | 16+17+18 | + | + | - | 140 |
| **2+3** | 7 | - | - | + | 34 |
| **7 (QR)** | 10+11 (QR) | - | - | + | 43 |
| **7 (QR)** | 16+17+18 (QR) |  |  |  | 90 |

**Figure 3:**
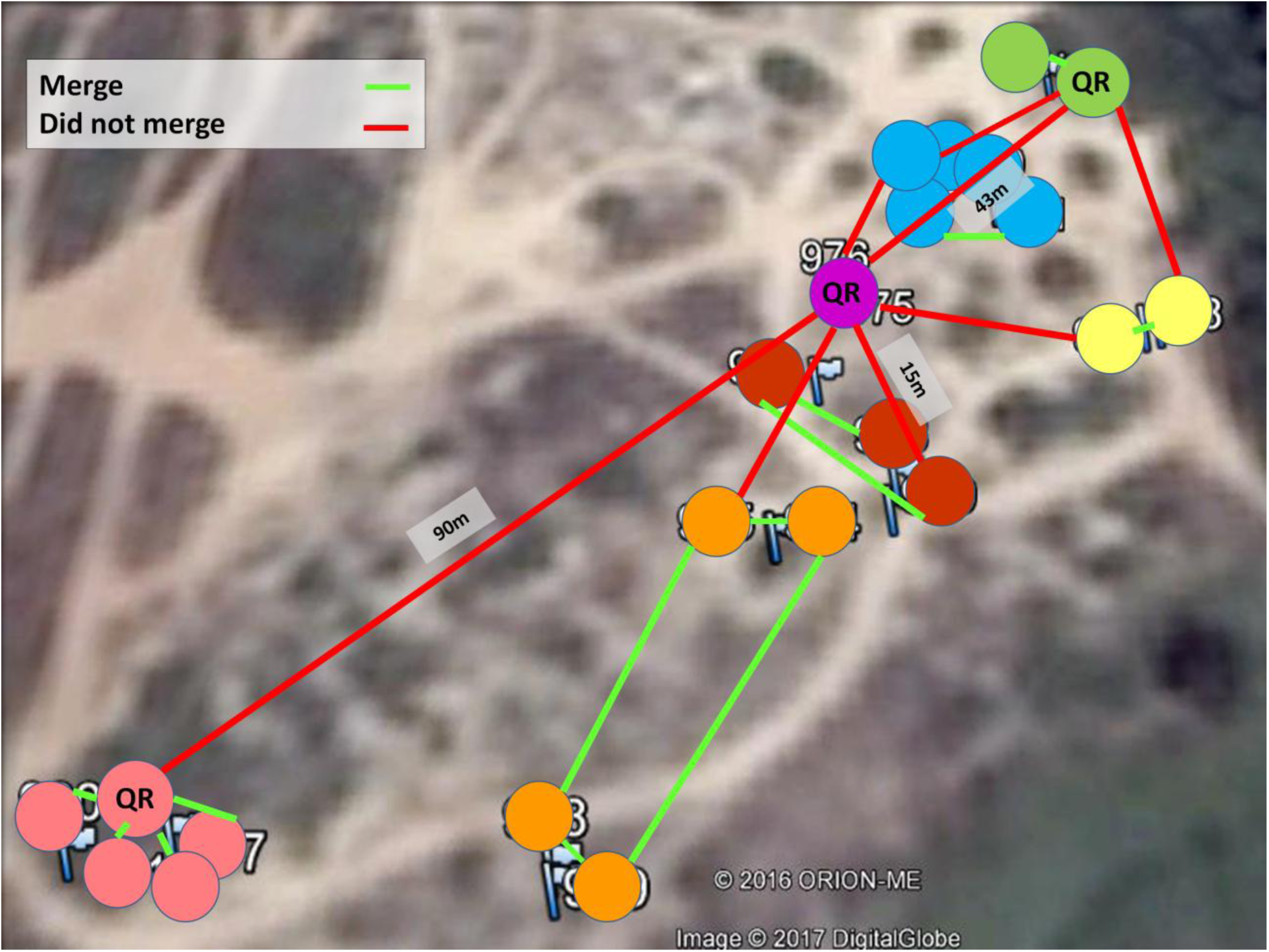
22 nests that were collected for the behavioral essays. Each circle indicates a physical nest, while each color indicates a different colony obtained by the merging assays. Queen-right nests are indicated by a large circle. Assays were performed between all colonies as indicated in Table 2. Nests that merged are indicated by a green line; nests that did not merge are indicated by a red line.

### Tel Baruch Population

#### Genetic analysis

Genetic analysis (N= 26 nests) using CytB identified representatives of all three mitotypes belonging to the ‘niger complex’ in the sampled plot *(niger, savignyi* and *drusus;* n= 6, 17 and 3 nests, respectively).

Microsatellite DNA analysis was performed on 548 workers of the Tel Baruch population (mean number of workers typed per nest ± SE= 21.08 ± 0.45, N= 26 nests) at 7 polymorphic loci. First, we analyzed all the nests in the population, irrespective of their mitotypes, assuming that they comprise a single population. The number of alleles ranged from five to 25. Mean heterozygosity was *Ho=* 0.69 (range: 0.4–0.87). The fixation index *Fit* was low, even though slightly higher than zero (mean ± SE jackknife= 0.02 +0.01) indicating that random mating predominates in the tested population. *Fst* was low at 0.12 while the within-nest relatedness was 0.23 (SE jackknife: 0.01). STRUCTURE indicated that the 26 nests analyzed consisted in 2, 3, or 6 colonies (Fig. 4A+B). Comparing colony distribution according to the above results (Fig. 4C) with that of the mitotype distribution (Fig. 4D) showed incongruence. Furthermore, PCA distribution of the genotypes showed that, overall, they were very similar throughout the population (Fig. 4E).

**Figure 4:**
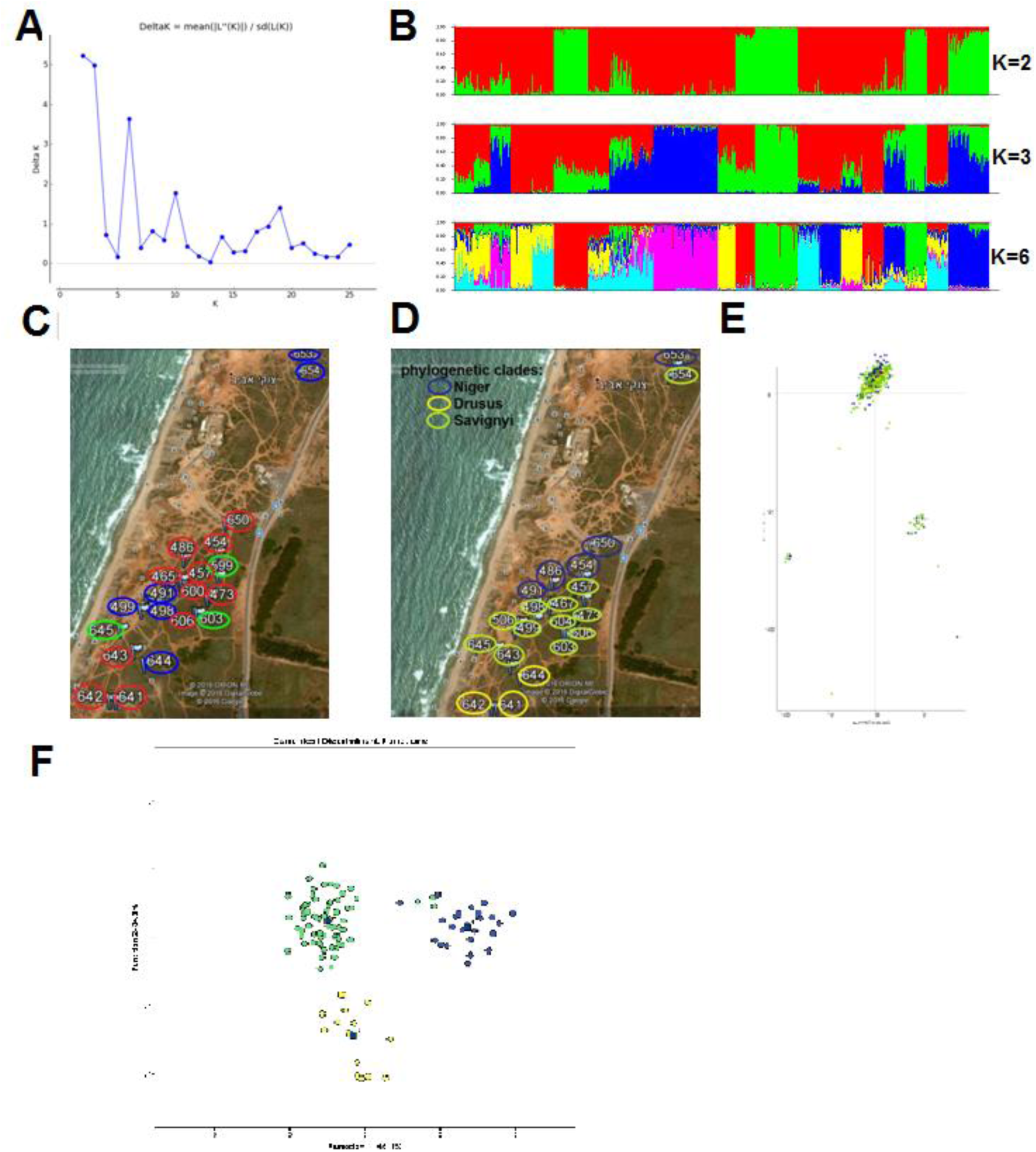
A) STRUCTURE harvester results for the Tel Baruch population indicating the best probable K is K=2, then K= 3 and K=6. B) The population STRUCTURE divisions according to the most probable K possibilities. C) STRUCTURE results for the Tel Baruch population of K=3 plotted on the map, the compression with the mitotypes plotted in (D) exemplifies that the STRUCTURE division is not related to the mitochondrial DNA. D) Plotted on the map of the Tel Baruch sampling site Cytochrome B phylogenetic clades. E) Principle Components Analysis of genotypes for Tel Baruch population colored according to the different mitotypes. F) Discriminant analysis results of CHC’s composition for the Tel Baruch population according to the three mitotypes.

Genetic analysis of the population structure considering the mitotypes separately gave a slightly different picture. Each mitotype had its distinct characteristics (Table 3); the *niger* mitotype had a very low *Fst* and very low within-nest relatedness, as predicted for polygyne colonies. The *drusus* and *savignyi* mitotypes, on the other hand, exhibited higher *Fst* and relatedness values, as predicted for monogyne colonies. However, perhaps due to the rather small sample size or the unique situation of this sympatric population, we obtained slightly different results to those for the other studied populations containing a single mitotype (data from: the Betzet population-only *drusus* mitotype in this paper, the Nitzanim population-only *niger* mitotype from Leniaud *et al*., 2011, and the Rishon and Arad populations-only *savignyi* mitotype from Leniaud *et al*., 2011; Saar *et al*., 2014; Table 3).

**Table 3:** Microsatellites analysis of the Tel Baruch population when divided according to the different mitotypes.

| Haplotype | Fit | Fit in a single mitotype population | Fst | Fst in a single mitotype population | Fis | Within nest Relatedness | Within nest Relatedness in a single mitotype population |
| --- | --- | --- | --- | --- | --- | --- | --- |
| **All nests/haplotypes**<br>**n=26** | 0.022±0.013 |  | 0.117±0.008 |  | 0.108±0.008 | 0.229±0.013 |  |
| **niger**<br>**n=6** | 0.051±0.02 | 0.023±0.008<sup>1</sup> | 0.059±0.005 | 0.005±0.003<sup>1</sup> | 0.008±0.019 | 0.112±0.009 | 0.02±0.007<sup>1</sup> |
| **drusus**<br>**n=3** | 0.027±0.033 | 0.09±0.06<sup>2</sup> | 0.092±0.008 | 0.19±0.015<sup>2</sup> | 0.131±0.029 | 0.19±0.013 | 0.35 ±0.02<sup>2</sup> |
| **savignyi**<br>**n=17** | 0.03±0.011 | 0.031<sup>3</sup> | 0.143±0.013 | 0.151<sup>3</sup> | 0.132±0.015 | 0.278±0.023 | 0.247<sup>3</sup> |

<sup>1</sup>Population is located in Nitzanim (Leniaud *et al.*, 2011)
<sup>2</sup> Population is located in Betzet (This paper)
<sup>3</sup> Population is located in Arad, Rishon and Palmachim (Leniaud *et al.*, 2011; Saar *et al.*, 2014)

#### Cuticular hydrocarbon analyses

For chemical analyses, we used head extracts of 76 ants belonging to 28 nests. We identified 74 compounds in the cuticular washes, of which 37 were used in the following analysis (supplementary Fig. S2B). Profile determination was constructed by calculating the relative amounts of each of the hydrocarbon components, and profile comparison was done by discriminant analysis, while grouping the ants according to the CytB mitotype. As can be seen in Fig. 4F, there was a clear discrimination of CHC profiles among the three mitotypes.

#### Behavioral assay

Two alternative hypotheses emerged from the genetics data regarding the social structure of the Tel Baruch population: a single, highly polymorphic species, as suggested by the nuclear gene analyses; or multiple species, as suggested by the mtDNA and CHC analyses. We therefore attempted to differentiate between the two hypotheses using behavioral assays comprising the possible merging of two distinct nests. Seven merging tests were performed using various pairs of nests (Fig. 5 and Table 4). First, we used nests of the *niger* mitotype located at distances between 100 and 500 meters from one another. In all cases the nests merged after a few hours, during which we observed both trophallaxis and mutual carrying between workers from the different nests. After 24 hours, the nests had fully merged, with the queens and brood having been moved into one of the nest boxes. All other pairing combinations, i.e., *drusus-niger, drusus-savignyi, savignyi - savignyi*, and *savignyi-niger*, resulted in aggressive behavior between the workers and none of these nests merged (Fig. 5).

**Figure 5:**
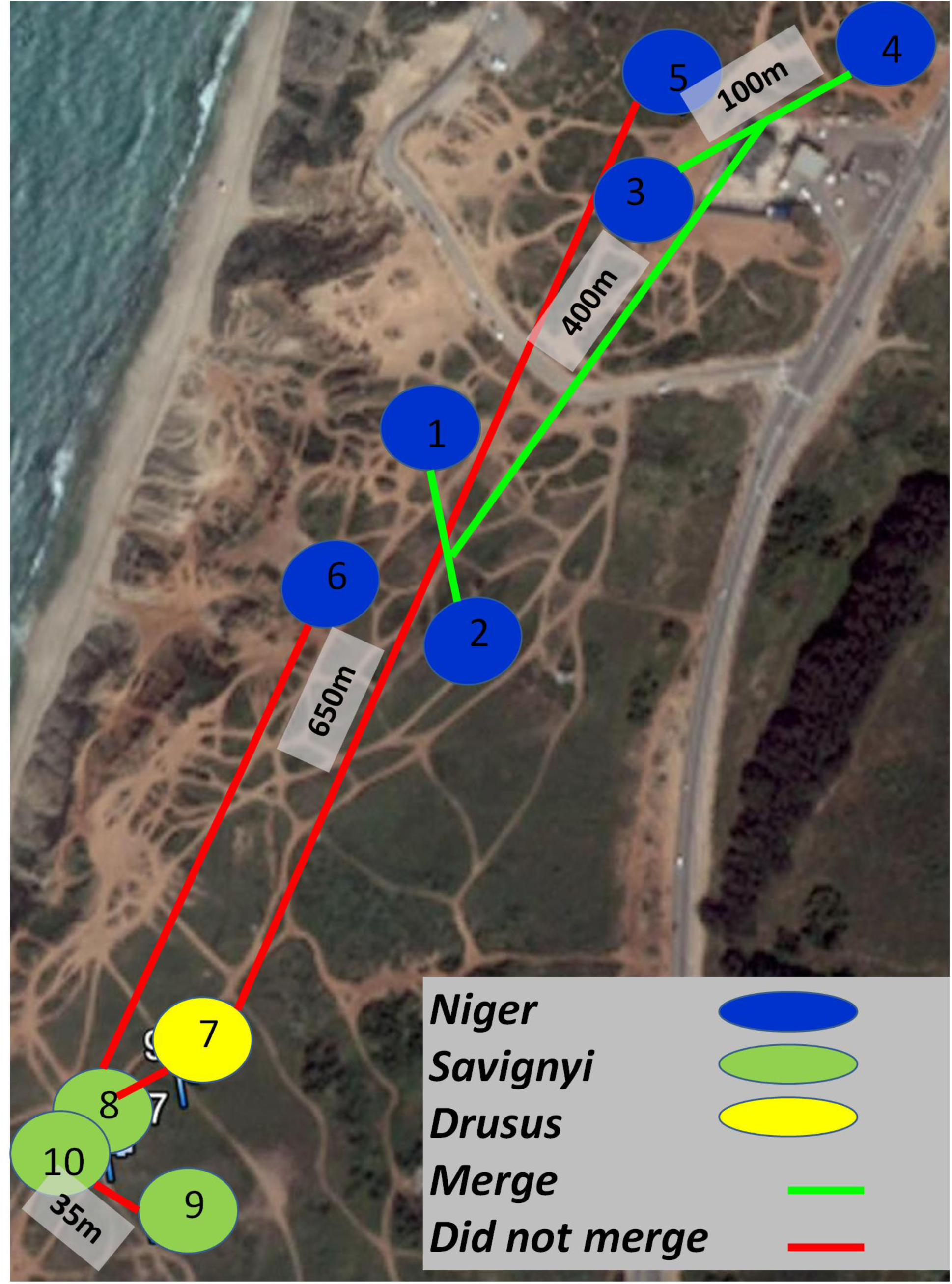
Location of nests used for the merging experiment. Nests are colored according to the mitotype: blue-*niger*, green-*savignyi*, and yellow-*drusus*. The lines drawn between nests indicate if the nests merged (green) or not (red).

**Table 4:** Merging assays of the Tel Baruch population

| **Nest A**<br>**(mitochondrial**<br>**haplotype)** | **Nest B**<br>**(mitochondrial**<br>**haplotype)** | **Merging** | **Between-**<br>**colony**<br>**trophallaxis** | **Between-**<br>**colony**<br>**aggression** | **Maximal distance**<br>**between colonies**<br>**(meters)** |
| --- | --- | --- | --- | --- | --- |
| **1 (*Niger*)** | 2 (*Niger*) | + | + | - | 100 |
| **3 (*Niger*)** | 4 (*Niger*) | + | + | - | 100 |
| **1+2 (*Niger*)** | 3+4 (*Niger*) | + | + | - | 400 |
| **5 (*Niger*)** | 7 (*Drusus*) | - | - | + | 650 |
| **6 (*Niger*)** | 8 (*Savignyi*) | - | - | + | 450 |
| **7 (*Drusus*)** | 8 (*Savignyi*) | - | - | + | 35 |
| **9 (*Savignyi*)** | 10 (*Savignyi*) | - | - | + | 35 |

#### Sperm analysis

To examine whether assortative mating between the different mitotypes occurs, we genotyped, using CytB, the sperm extracted from the spermatheca of ten queens of the *niger* mitotype. Nine out of the ten sperm samples matched the *savignyi* mitotype and one matched the *niger* mitotype (Fig. 6). In some of the sequences there appeared to be more than one possibility of a nucleotide in certain positions, possibly indicating multiple sequences/mitotypes. This suggests that multiple males of multiple mitotypes inseminated a specific queen.

**Figure 6:**
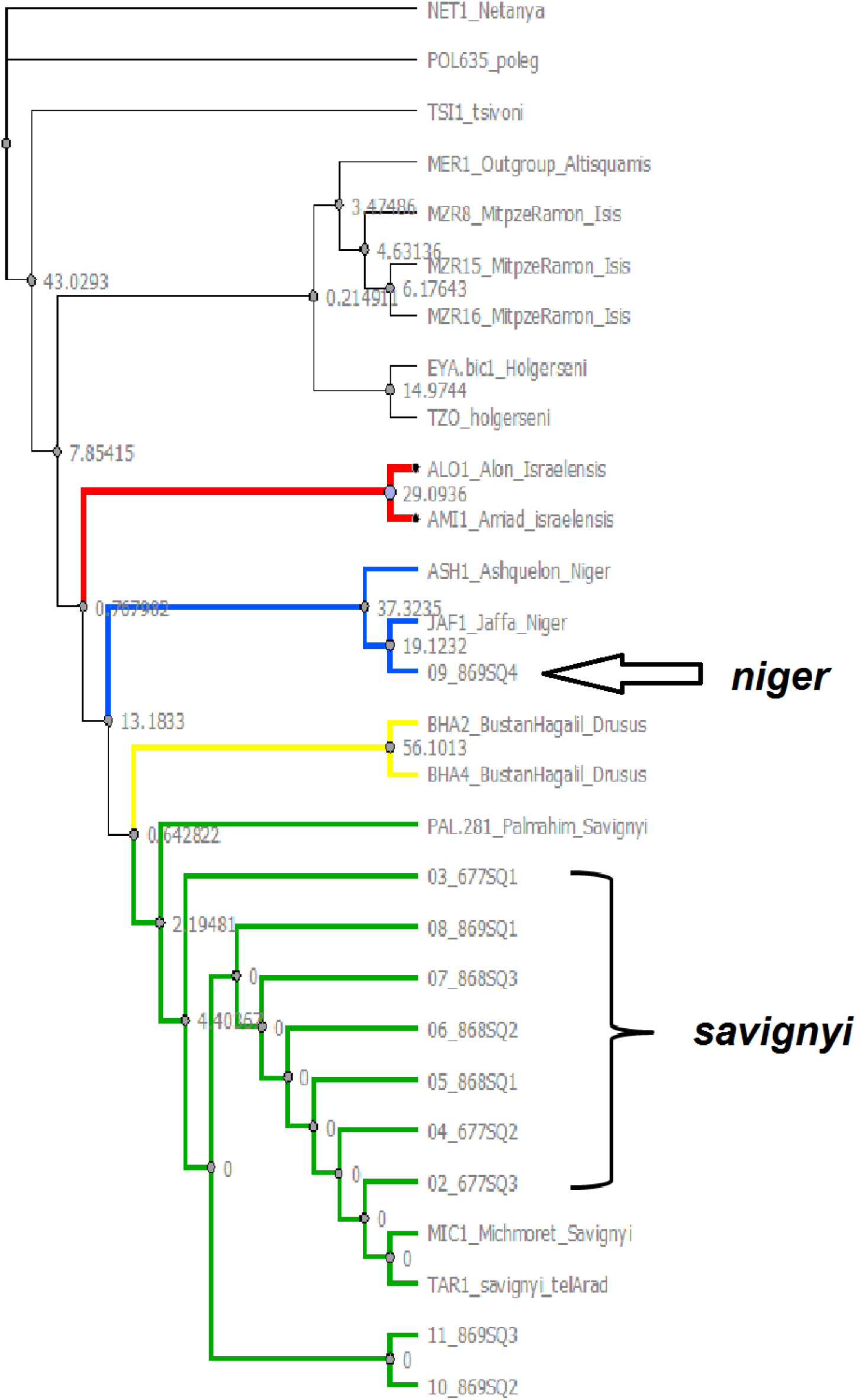
Bayesian inference tree CytB haplotypes of sperm content from *niger* queens. Bayesian probabilities/bootstrap values (from Ugene) are given to estimate branch support. The trees are rooted using the haplotype sequences from Eyer *et al*. (2017).

## Discussion

A recent phylogeographic study of species delimitation in the *bicolor* group of the genus *Cataglyphis* in Israel (Eyer *et al*., 2017) pointed out discrepancies between mitochondrial DNA and nuclear DNA sequences within the putative species *C. niger, C. savignyi*, and *C. drusus*, i.e., the *C. niger* complex. The three mitotypes also differ in their social structure. The *savignyi* mitotype is monogyne demonstrating either monodomous (Leniaud *et al*., 2011) or polydomous (Saar et al., 2014; mistakenly identified as *niger* when published) nesting behavior; whereas the *niger* mitotype is polygyne and is organized as a supercolony (Leniaud *et al*., 2011). The social and population structure of the *drusus* mitotype has not previously been investigated. Furthermore, this study revealed that all three mitotypes occur sympatrically in the Tel Baruch site. Therefore, disentangling the species identity of this complex is expected to provide insight not only into the process of sympatric speciation, but also into the evolution of advanced social structures in ants.

First, we sought to rule out the possibility that the amplicons of the nuclear genes employed in Eyer et al. (2017) were not sensitive enough to detect differences between the three mitotype. We therefore conducted RAD sequencing on representatives of the above complex, along with samples from most of the locations described in Eyer et al. (2017), comprising other species, namely *C. holgerseni, C. isis, C. israelensis* and *C. nodus* (collected in Turkey). The RAD results were compatible with the nDNA data of the Eyer et al., (2017) study, and ruling out the possibility of errors due to the limited sequence information for the study. *STRUCTURE* results confirmed the existence of five bicolor species in Israel, and most importantly confirmed the existence of gene flow between the three mitotypes of the *C. niger* complex (i.e. *niger, savignyi*, and *drusus)*, suggesting that they constitute a single species. The results also indicate hybrid zones between *C. israelensis* and the *savignyi* mitotype of the *C. niger* complex.

The case of the *C. niger complex* is interesting and somewhat enigmatic. The three mitotypes have a peculiar distribution along the coastal plains of Israel (Eyer *et al*., 2017). The northern part, from the Lebanese border to Mount Carmel, is inhabited exclusively by the *drusus* mitotype. To the south of Mount Carmel, the *savignyi* mitotype predominates, except for a patchy distribution of another species, *C. israelensis* (Ionesco and Eyer, 2016; Eyer *et al*., 2017). Curiously, at the border of their distribution, we found hybrids between *C. israelensis* and the *savignyi* mitotype of the *C. niger* complex. *The savignyi* mitotype has the largest distribution. It occupies a large part of the sandy areas along the coast and it dominates in areas of heavier soil, extending to the more desert areas of Israel. The *niger* mitotype, in contrast, is present only in the central coastal plain, where it has a patchy occurrence. Thus, at least in a large proportion of their distribution, although the *drusus* and *savignyi* mitotypes are allopatric, the nuclear sequencing data indicates free gene flow. This raises the question of whether we are encountering 3 incipient species that demonstrates rapid mitochondrial evolution but slower nuclear evolution; or a single species with long-range dispersal of sexuals (mostly through males because *Cataglyphis* females are generally known to have lower flying capability; Peeters and Aron, 2017), to account for the apparent gene flow among the highly allopatric populations.

The sympatric cooccurrence of all three mitotypes in the Tel Baruch population enables us to investigate the differences among the complex mitotypes within a natural setting without a confounding ecological variation. The microsatellite analysis reveals an *Fst* value of 0.12 across all the nests of the Tel-Baruch population, suggesting some level of diversification in the highly polymorphic markers (Wondji, Simard and Fontenille, 2002; Leppänen *et al*., 2015). However, this estimate might not be accurate because some nests belong to the same colony. As reported previously, the *niger* mitotype is polygyne and supercolonial (Leniaud *et al*., 2011). Here, we further established that this is an exclusively *niger* mitotype characteristic. Both *savignyi* and *drusus* at the study sites were monogyne and facultatively polydomous. The three mitotypes inhabiting the same plot were also distinctively different in their CHC composition. CHC’s are the main mechanism through which ants maintain nest insularity (Soroker *et al*., 1994; Hefetz, 2007; Kidokoro-Kobayashi *et al*., 2012), thus making it highly important in such a diverse population. Within each of the mitotypes, differences in CHC profiles were rather small, consistent with populations exhibiting limited dispersal. Thus, the question of whether we have 3 incipient species (as indicated by morphology (Agosti, 1990), mitochondrial DNA, chemistry, and behavioral data), or a single species (as indicated by the nDNA genotyping) that is highly polymorphic (morphologically, chemically and to a certain degree genetically), remains open. While this does not pose a problem regarding mitotype stability at the site (being maternally inherited), how the social structure (i.e., polygyny and supercoloniality) is maintained remains elusive.

According to the incipient species hypothesis, we expected assortative mating according to the different mitotypes. However, sperm analysis of mated gynes of the *niger* mitotype revealed a high presence of the *savignyi* mitotype (matching its abundance at the site), indicating the absence of mitotype-associated assortative mating. Such non-assortative mating can be the result of either recent admixture of the 3 species due to anthropogenic-associated geographic barrier destruction, leading the 3 species to merge into a single species, or supporting the single species hypothesis having 3 mitochondrial lineages, each of which associated with different social structure. Cross-mating between lineages within a species has been described in *Pogonomyrmex*, resulting in caste differentiation: eggs fertilized by males of the queen’s lineage develop into new queens, while eggs fertilized by males of the opposite lineage develop into sterile workers (Schwander, Keller and Cahan, 2007). Another report of cross-mating between two social forms is that for *Solenopsis invicta* (Shoemaker and Ross, 1996; Ross and Shoemaker, 1997), in which polygyne queens mate with males from monogyne colonies.. In *Cataglyphis*, several species of the closely-related *C. altisquamis* group exhibit cross-mating between two hybridogenetic lineages, which facilitates worker production and genetic diversity (C. *velox, C. altisquamis, C. hispanica* and *C. mauritanica* Leniaud *et al*., 2012; Aron, Mardulyn and Leniaud, 2016; Boulay *et al*., 2017a). Here we showed that the polygyne *niger* queens partake in nuptial flights, which enables mating with males of the *savignyi* mitotype. However, since within-colony genetic diversity is satisfied in the case of non-related co-inhabiting queens (Reiner Brodetzki *et al*., 2018), it is not clear whether there is a genetic necessity for cross-mitotype mating, paving the way for the evolution of assortative mating among mitotypes. In other *Cataglyphis* polygyne species, gyne morphology reveals the loss of wing muscles (Peeters and Aron, 2017), confining the queens to mating near or within the nest. Such a change in morphology would lead to mating place separation and sympatric speciation. Another possible mechanism that could explain the social polymorphism exhibited by the *niger* complex, if considering the complex as a single species, is the occurrence of a non-combining ‘social chromosome’, as described in two other ant species, *Solenopsis invicta* and *Formica selysi* (Wang *et al*., 2013; Purcell *et al*., 2014). The independent evolution of a ‘social-chromosome’ mechanism may suggest that such a mechanism may also have evolved in *Cataglyphis*. However, the apparent differences in the mitochondrial DNA may indicate a maternal inheritance of these social traits as well, which raises the possibility of alternative explanations, not previously described in ants, such as a maternal genetic imprinting on social traits.

When considering possible mechanisms and road maps for sympatric speciation, we assume traits evolving in a single population in opposite directions. These traits should have major life-history effects that would drive for assortative mating, stopping gene flow and promoting speciation. Social behavior and recognition cues are such traits. The *C. niger complex* exemplifies a population in which some major traits have diverged while others have not, raising the possibility of a sympatric speciation continuum based on a differentiation in sociality.

## Acknowledgments

This research was funded by the Israel Science foundation (ISF grant # 844/13 to AH). The research was also supported by the Naomi Prawer Kadar Foundation through the Tel Aviv University GRTF Program (travel grant to TRB). We would like to thank Raya Zeltzer, Maayan Drin, and PA Eyer for their help with sample collection; Guy Sobol, Yafit Brenner, and Tovit Simon, for technical assistance and data process; Laurent Grumiau and Hugo Darras for help with genetic data; and Naomi Paz for editing.

## Supplementary figures

**Figure S1:**
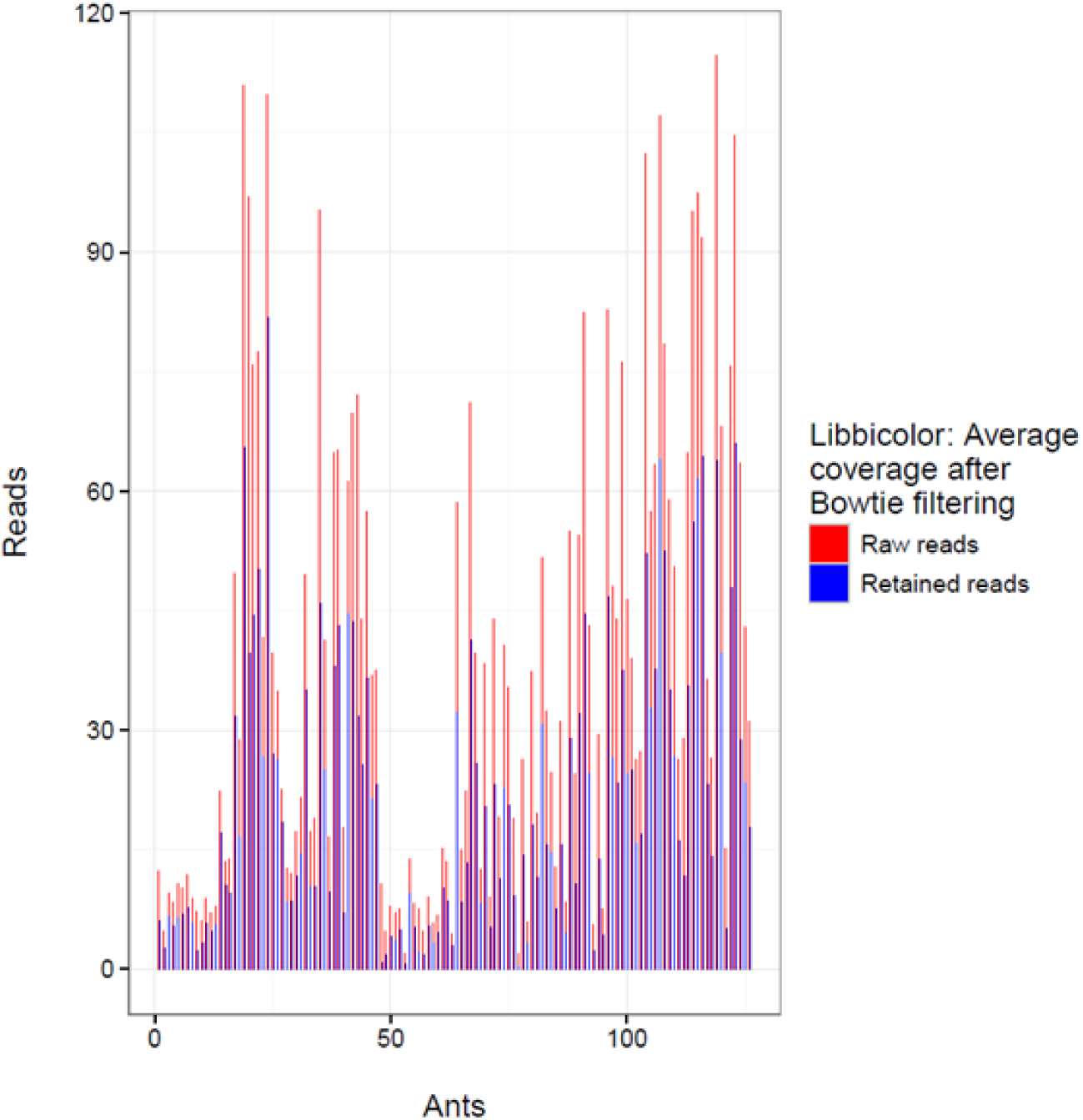
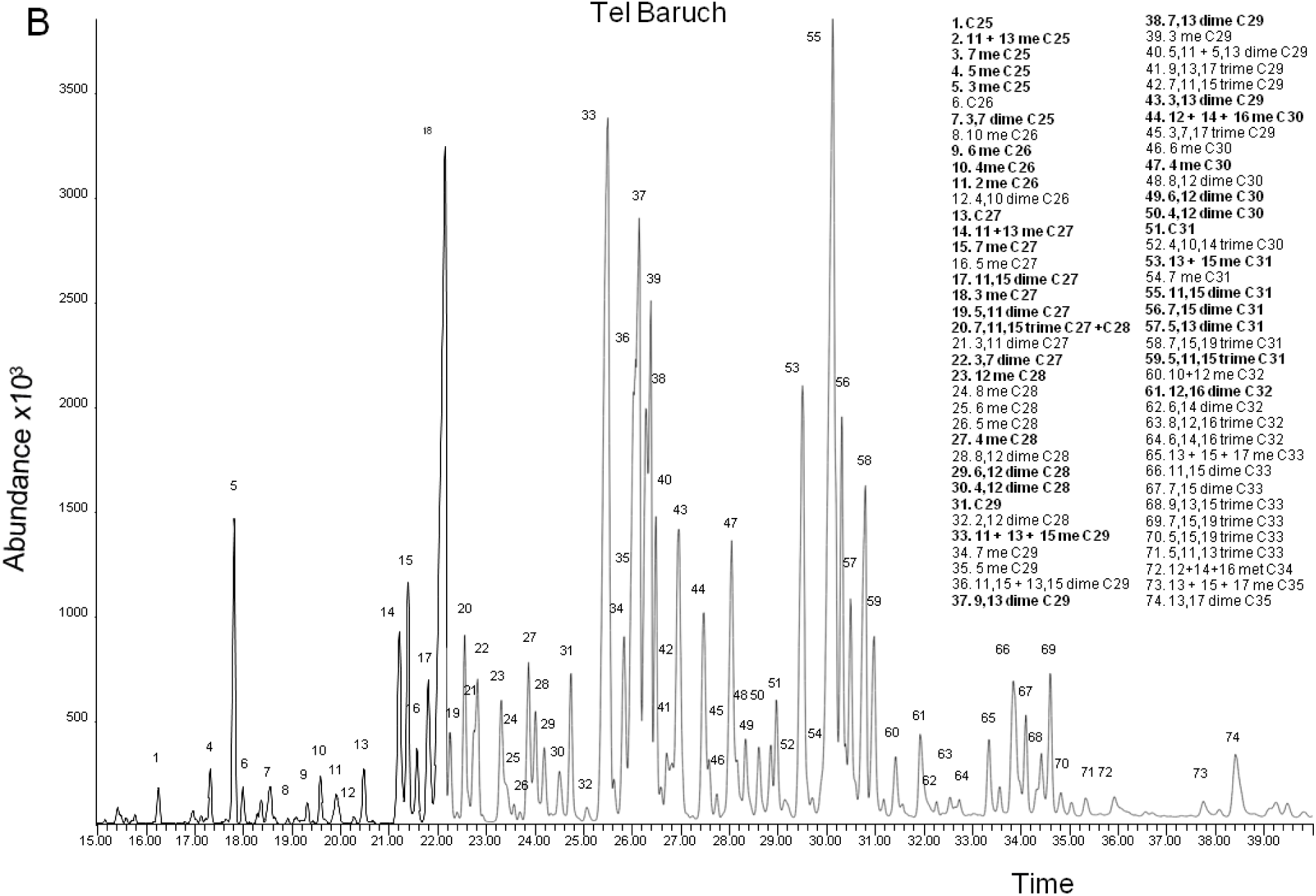
comparison of reads coverage of each ant at the stage of raw reads (red) and after the alignment based filtering (blue). Notice that the initial coverage is based on averaging the number of reads over the known number of restriction sites, while the final coverage is based on actual mapping of the reads to the reference genome of *C. drusus* and is therefore more accurate.

**Figure S2:**
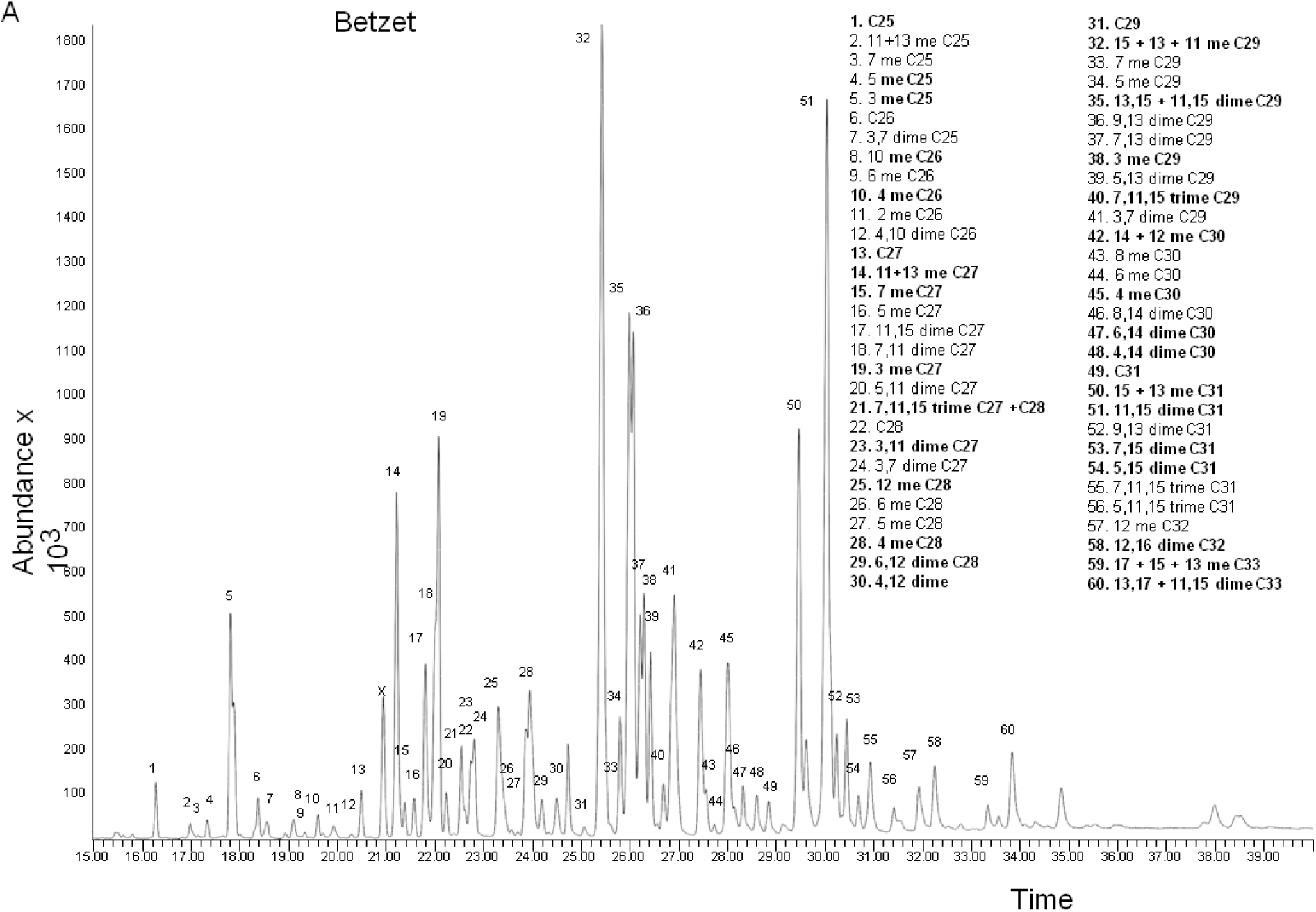
A) Typical cuticular hydrocarbon profile of the *drusus* mitotype from Betzet. The chromatogram shows only the 60 identified compounds (numbered peaks), of which 35 hydrocarbons were used in the analysis (marked in bold). B) The cuticular hydrocarbon profile from the Tel Baruch population CHC’s used for the analysis is marked in bold (44 out of 74 identified).

## Supplementary Tables

**Table 3.S1:**
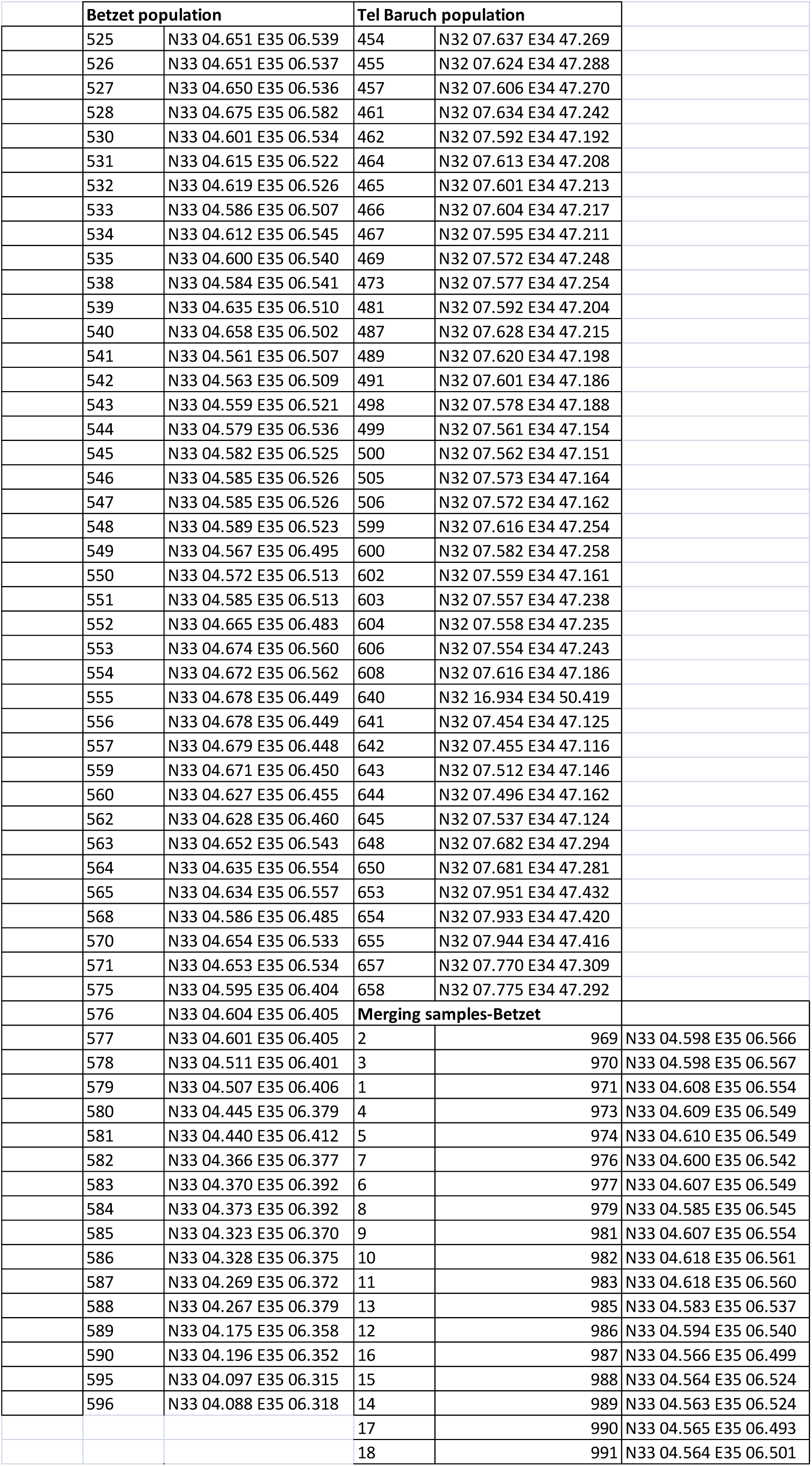
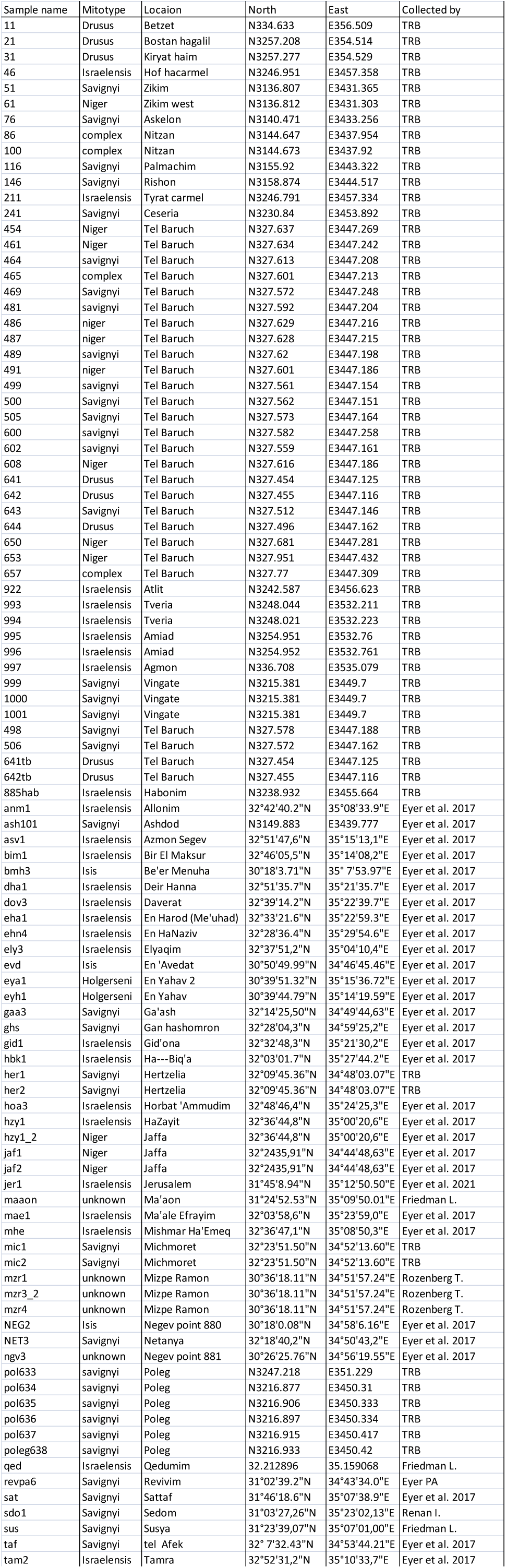
GPS coordinates of samples used in the study.

**Table S2:** The number of nests used in each of the analysis.

| **Location** | **QL nests** | **QR nests** | **Nests in merging experiment** | **Nests in microsat analysis** | **Nests in CHC analysis** |
| --- | --- | --- | --- | --- | --- |
| **Betzet** | 76 | 5 | 22 | 57 | 49 |
| **Tel Baruch** | 39 | 21 | 10 | 26 | 28 |

**Table S3:**
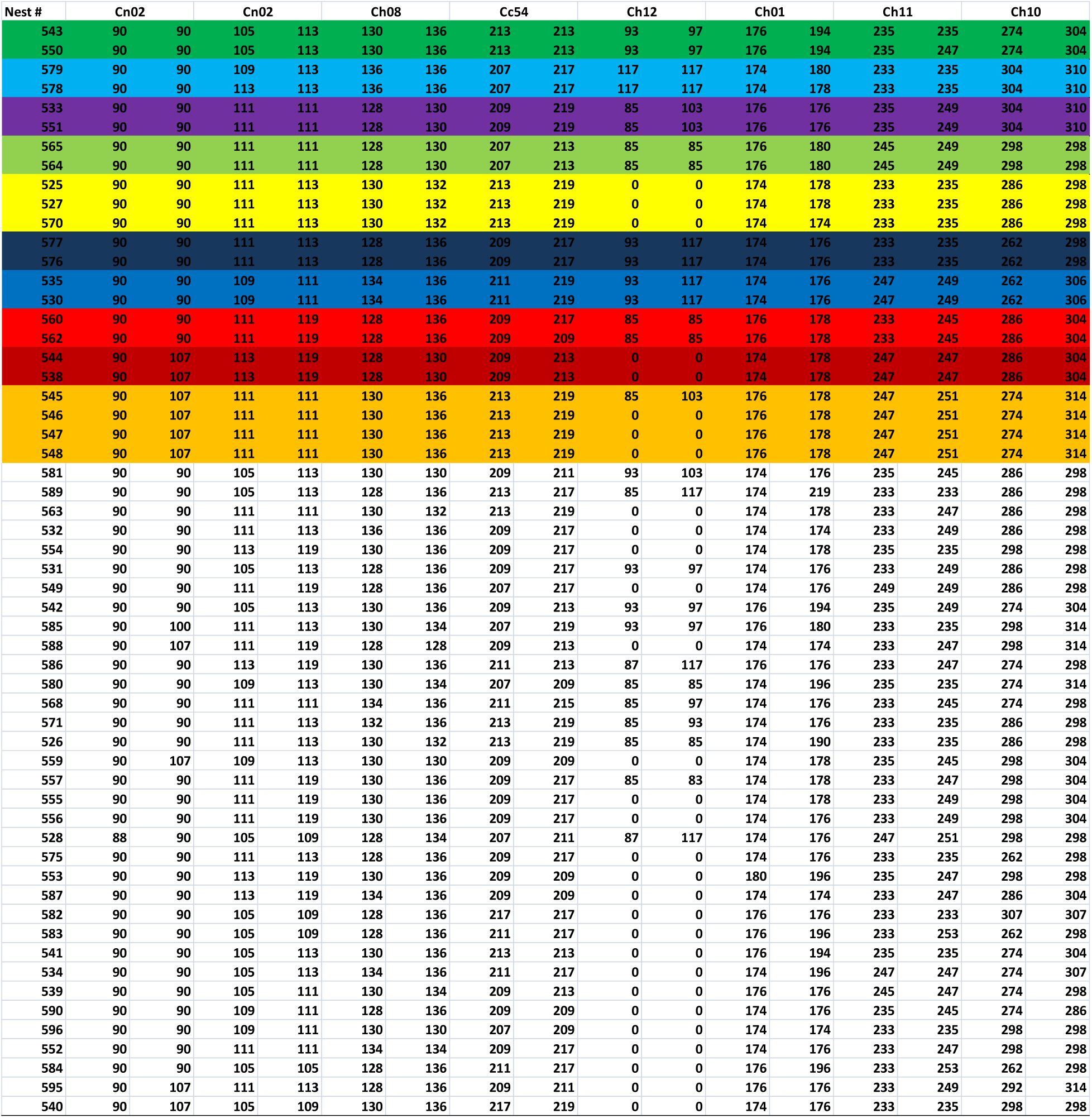
Assumed matrilines according to COLONY in the Betzet population.

